# Non-Redundant Roles in Sister Chromatid Cohesion of the DNA Helicase DDX11 and the SMC3 Acetyl Transferases ESCO1/2

**DOI:** 10.1101/704635

**Authors:** Atiq Faramarz, Jesper A. Balk, Anneke B. Oostra, Cherien A. Ghandour, Martin A. Rooimans, Rob M.F. Wolthuis, Job de Lange

## Abstract

In a process linked to DNA replication, duplicated chromosomes are co-entrapped into large, circular cohesin complexes and functional sister chromatid cohesion (SCC) is established by acetylation of the SMC3 cohesin subunit. Several rare human developmental syndromes are characterized by defective SCC. Roberts Syndrome (RBS) is caused by mutations in the SMC3 acetyl transferase ESCO2, whereas mutations in the DNA helicase DDX11 lead to Warsaw Breakage Syndrome (WABS). We found that WABS-derived cells predominantly rely on ESCO2, not ESCO1, for residual SCC, growth and survival. Reciprocally, RBS-derived cells depend on DDX11 to maintain low levels of SCC. Synthetic lethality between DDX11 and ESCO2 correlated with a prolonged delay in mitosis, and was rescued by knock down of the cohesion remover WAPL. Rescue experiments using mouse or human cDNAs revealed that DDX11, ESCO1 and ESCO2 act on differential aspects of DNA replication-coupled SCC establishment. Importantly, DDX11 is required for normal DNA replication fork speed without clearly affecting SMC3 acetylation. We propose that DDX11 and ESCO2 control spatially separated fractions of cohesin, with supportive roles for DDX11 in replication-associated cohesion establishment, and for ESCO2 in cohesin complexes located around centromeres, respectively.

## Introduction

Sister chromatid cohesion (SCC) is mediated by cohesin, a presumed DNA-entrapping ring formed by structural maintenance of chromosome 1 and 3 (SMC1 and SMC3), RAD21 and SA1/2. The loader complex MAU2-NIPBL loads DNA into cohesin rings [1–3], whereas it can be released by the cohesin remover WAPL [4]. During DNA replication, stable cohesion is established in a process involving SMC3 acetylation by ESCO1 and ESCO2, which leads to the recruitment of Sororin and subsequent inhibition of WAPL activity [5–7]. The resulting SCC facilitates proper chromosome biorientation and equal distribution of genetic material during mitosis. Prior to chromatid separation in anaphase, cohesin needs to be removed, which happens in two rounds and via two distinct pathways [8, 9]. First, the prophase pathway promotes removal of cohesins from chromosome arms by WAPL, in a process involving multiple phosphorylations that restore WAPL activity [10]. Centromere cohesins are protected from the prophase pathway by SGOL1, which recruits the PP2A phosphatase to the centromeres [9, 11, 12]. In a separate step that occurs at the metaphase-to-anaphase transition, the remaining centromeric cohesins are removed by the protease Separase, which is activated by the Anaphase-Promoting Complex/Cyclosome (APC/C) and cleaves the RAD21 subunit [13]. In addition to its role in sister chromatid cohesion, the capacity of cohesin to entrap DNA also allows it to regulate gene transcription [14–16] and promote ribosome biogenesis [17–19].

Mutations in cohesin components or regulators result in a cluster of syndromes called cohesinopathies, characterized by diverse clinical abnormalities including growth retardation, intellectual disability, microcephaly and congenital abnormalities. Four cohesinopathies have been described thus far. Cornelia de Lange syndrome (CdLS) results from autosomal dominant or X-linked mutations in NIPBL, SMC1A, SMC3, RAD21 or HDAC8 [20–26]. Roberts Syndrome (RBS, also called SC phocomelia syndrome) is caused by bi-allelic mutations in ESCO2 [27]. Warsaw breakage syndrome (WABS) results from bi-allelic mutations in the DNA helicase DDX11 [28]. Chronic Atrial and Intestinal Dysrhythmia (CAID) syndrome was described in a patient with homozygous missense mutations in SGOL1 [29]. CdLS cells exhibit no defects in SCC [30], and the clinical symptoms are thought to originate from deregulated gene transcription (reviewed in [31–33]). By contrast, metaphases derived from RBS, WABS and CAID patient cells show severe cohesion loss [27–29]. The clinical symptoms of these syndromes are likely to originate from a combination of transcriptional defects and reduced progenitor cell proliferation.

ESCO1 and ESCO2, the vertebrate orthologs of yeast Eco1, share a conserved C-terminus that contains a zinc finger motif and an acetyltransferase domain, whereas no similarity is found in the N-terminus [34]. ESCO2 deficiency is embryonic lethal in mice, indicating that ESCO2 functions non-redundantly with ESCO1 [35]. RBS patient derived cells show defective centromere cohesion [36], in line with the observation that ESCO2 localizes to pericentric heterochromatin [35]. ESCO2 expression peaks during S-phase and is subsequently reduced by proteasomal degradation [35, 36], indicating that its prime function is to mediate centromeric sister chromatid cohesion in the context of DNA replication. In budding yeast, Eco1 is reported to be recruited to the replication fork by replication factor PCNA [37] and decreased SMC3 acetylation in human cells reduces replication fork speed [38]. The authors proposed that the primary role of SMC3 acetylation would be to switch cohesin from a conformation that obstructs replication forks to a more open structure that allows fork progression [38]. Unlike ESCO2, ESCO1 is expressed during the whole cell cycle and has been reported to acetylate SMC3 independent of replication, suggesting that ESCO1 also regulates non-canonical roles of cohesin [39]. Nevertheless, ESCO1 knockdown was also found to cause cohesion loss in Hela cells [34, 39] and DT40 chicken cells [40]. A different study reported no effect of ESCO1 knockdown or CRISPR mediated knockout on cohesion in Hela cells, and the authors proposed a model in which ESCO1 facilitates structural cohesion rather than replicative cohesion, thereby indirectly reinforcing cohesion that was established by ESCO2 [41].

DDX11 belongs to a group of ATP-dependent, super-family 2 (SF2) DNA helicases with an iron-sulfur cluster (Fe-S) domain [42]. It is specialized in unwinding certain DNA structures that contain a 5’-single stranded region, including forked duplexes, 5’-flap duplexes and anti-parallel G-quadruplexes (reviewed in [43]). This may be particularly relevant in the context of a DNA replication fork, where potentially long stretches of single stranded DNA can form secondary structures. Indeed, DDX11 and its yeast ortholog Chl1 have been shown to interact with multiple replication factors, such as the sliding clamp PCNA, its loader Replication Factor C complex (RFC), the 5’-flap endonuclease FEN1 and CTF4, which couples the MCM helicase to DNA polymerases [44–46]. How DDX11 deficiency results in cohesion loss is not entirely understood. The resolution of complex secondary DNA structures that are formed particularly in the lagging strand may be required for successful sister chromatid entrapment, but more direct, helicase-independent roles in cohesin loading have also been proposed [47–49]. Interestingly, Eco1 and Chl1 genetically interact [50–52] and synthetic lethality between DDX11 and ESCO2 was also reported in chicken DT40 cells [53].

Here we report synthetic lethality between ESCO2 and DDX11 in different human cell lines. Lethality is accompanied by aggravated cohesion loss in metaphase spreads. Reinforcing arm cohesion by WAPL knockdown rescues synthetic lethality, indicating that lethality results from too severe loss of SCC. ESCO1 and ESCO2 appear to have both overlapping and non-overlapping roles in SCC, which are conserved between mice and men. Finally, we show that DDX11 deficient cells have reduced replication fork speed, comparable to ESCO2 deficient cells. We conclude that DDX11 and ESCO2 have functions in SCC that are both spatially and mechanistically distinct. Whereas ESCO2-mediated SMC3 acetylation stabilizes cohesion particularly in centromeric regions, we speculate that DDX11 facilitates both DNA replication and cohesin loading by resolving complex secondary structures in difficult-to-replicate regions.

## Results

### Synthetic lethality of DDX11 and ESCO2 is conserved in different human cell lines

We previously generated a unique isogenic cell line pair by functionally correcting fibroblasts derived from a WABS patient [54] and used these in a genome-wide siRNA screen to search for genes that are synthetically lethal with mutant DDX11 [55]. ESCO2 was one of the strongest hits as validated by deconvoluting the siRNA pool (Figure S1). This confirms that the synthetic lethality between DDX11 and ESCO2, observed between yeast Chl1 and Eco1 [50–52] and chicken DDX11 and ESCO2 [53], is conserved in patient-derived cells. We used the same WABS cell line and its complemented isogenic pair to further validate these findings and also to examine the role of ESCO1 in this context. As expected, siRNA mediated ESCO2 knockdown strongly reduced the viability of WABS cells, but not of WABS+DDX11 cells (Figure 1A). This lethality correlated with increased levels of caspase-dependent PARP cleavage, reflecting apoptosis induction (Figure 1B). By contrast, knockdown of ESCO1 did not significantly affect cell growth in these cells, despite a substantial reduction of acetylated SMC3 which is the main target of ESCO1/ESCO2 acetyltransferases (Figure S2). In addition to these effects on viability, we observed severely aggravated cohesion defects in WABS cells upon knockdown of ESCO2, whereas ESCO1 knockdown had a moderate effect (Figure 1C). Reversely, DDX11 knockdown exhibited a stronger effect on cohesion in RBS cells than in RBS cells corrected with ESCO2 cDNA (Figure 1D, E).

**Figure 1:**
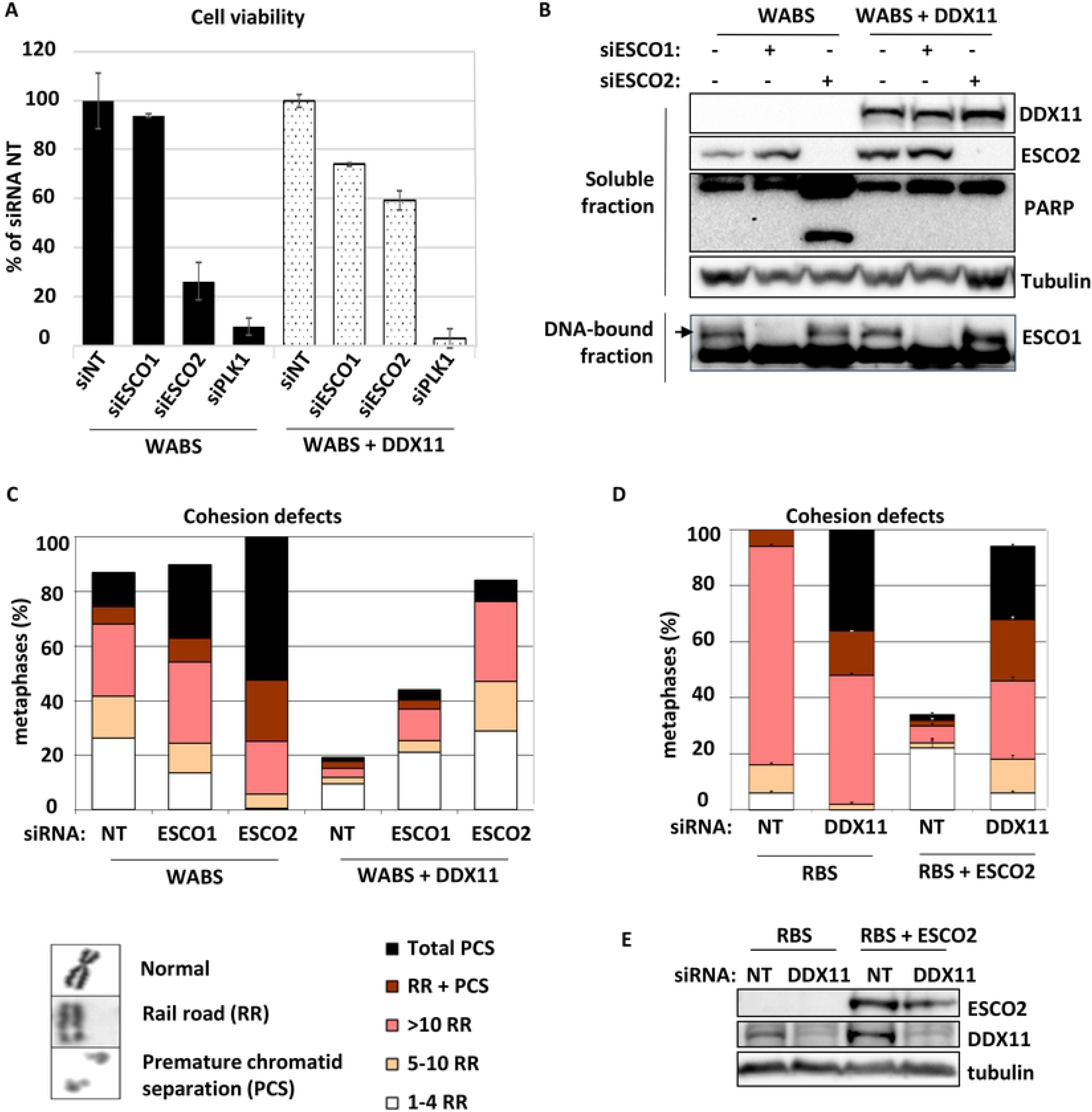
Synthetic lethality of DDX11 and ESCO2 is conserved in fibroblasts of WABS and RBS patients. (A) WABS fibroblasts and corrected cells were transfected with the indicated siRNAs and cell viability was analyzed after four days, using a cell-titer blue assay. (B) Cells were transfected with indicated siRNAs and analyzed by western blot after three days. (C) Cohesion defects were quantified in metaphases spreads of cells transfected as in B. Examples of metaphase chromosomes with normal and railroad (RR) appearance, as well as premature chromatid separation (PCS), are shown. (D, E) RBS fibroblasts and corrected cells were transfected with the indicated siRNAs and assessed after three days by cohesion defect analysis and western blot.

For additional validation, we used CRISPR-Cas9 to create RPE1-TP53KO cells, and subsequently generated DDX11KO and ESCO2KO in this genetic background. Knockdown of ESCO2 specifically inhibited growth of RPE1-TP53KO-DDX11KO cells (Figure 2A) and caused increased levels of PCS (Figure 2B), showing synthetic lethality cannot be rescued by TP53 loss. Similarly, cohesion defects were severely aggravated upon DDX11 knockdown in RPE1-TP53-ESCO2KO cells (Figure 2C).

**Figure 2:**
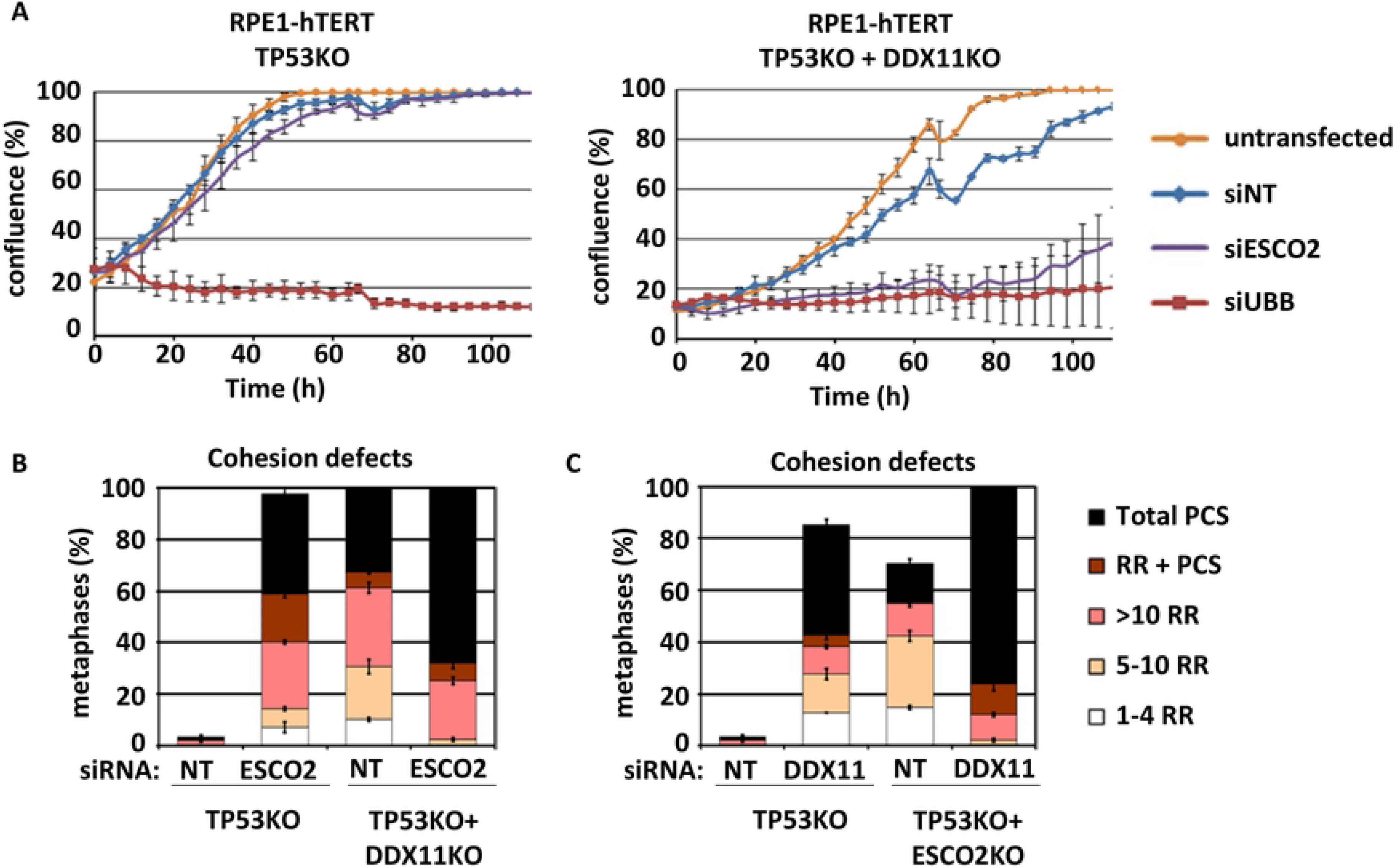
Synthetic lethality of DDX11 and ESCO2 in RPE1 cells. (A) CRISPR-Cas9 was used to knockout TP53 and DDX11 in RPE1-hTERT cells. The resulting isogenic cell lines were transfected with indicated siRNAs and proliferation was assessed using the IncuCyte. (B) Cells were transfected as in A and analyzed for cohesion defects. (C) CRISPR-Cas9 was used to disrupt the ESCO2 gene in RPE1-hTERT-TP53KO cells. Cells were transfected with indicated siRNAs and analyzed for cohesion defects

We further analyzed the effect of ESCO2 knockdown in WABS fibroblasts using flow cytometry, and found a clear induction of cells with a 4N DNA content (Figure 3A-C). This includes both p-Histone H3 positive (mitosis marker) and p-Histone H3 negative cells (G2 cells, 4N G1 cells). The dotplots (Figure 3B) also display the specific appearance of a new fraction with 4N DNA content that stains negative for p-Histone H3 (day 3), possibly dying cells. In addition, some polyploid (>4N) cells appear at day 3. These findings probably reflect an extended metaphase duration via reactivation of the spindle assembly checkpoint that follows from premature chromatid separation, as previously reported by us and others [55–57]. Eventually these arrested cells die in mitosis (as monitored by an increase in sub-G1 fraction), or may slip out of mitosis without cytokinesis. Cells that slip out of mitosis may subsequently continue to replicate leading to polyploidy. The increased 4N fraction could in part also reflect a G2 arrest, resulting from replicative stress or reduced DNA repair capacity.

**Figure 3:**
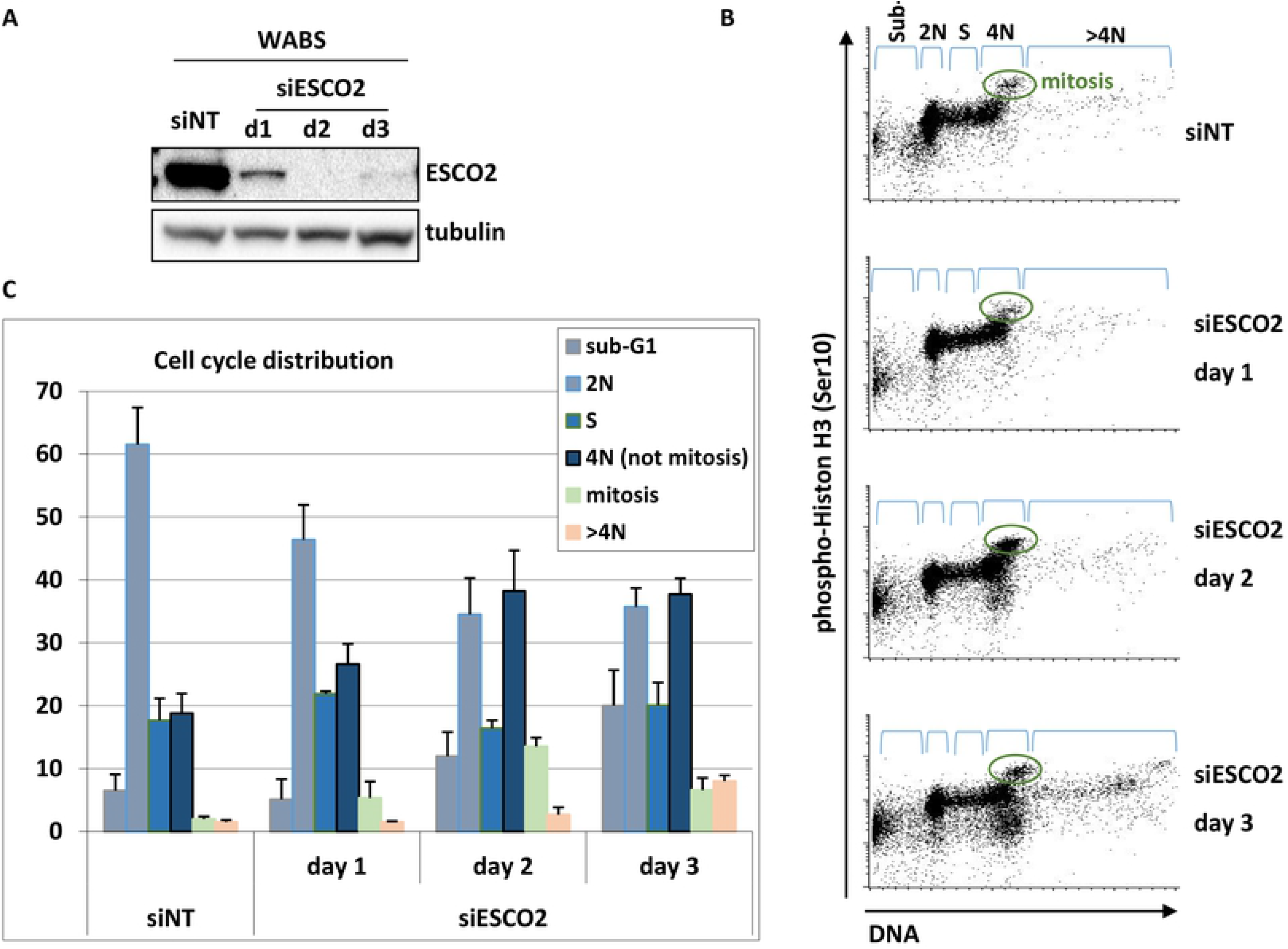
Induction of 4N fractions upon ESCO2 knockdown in WABS fibroblasts. WABS cells were transfected with the siESCO2, harvested at the indicated time points and analyzed by western blot (A) and flow cytometry (B). (C) Quantification of two independent experiments.

### Synthetic lethality of DDX11 and ESCO2 is rescued by WAPL knockdown

Next, we tested whether the SCC defects caused by single or double depletion of DDX11 and ESCO2 can be rescued by WAPL knockdown. WAPL plays a key role in removing cohesin rings from chromosome arms during the prophase pathway, so WAPL knockdown specifically causes hyper-cohesion on chromosome arms during metaphase. In agreement with specific roles for WAPL in arm cohesion, separated from a role for ESCO2 in centromere cohesion, the majority of railroads observed in RBS cells could not be rescued by WAPL knockdown (Figure 4A,B). However, WAPL knockdown prevented PCS, and, interestingly, rescued both synthetic lethality and PCS by ESCO2 knockdown in WABS cells (Figure 4C-E). The remarkable increase in railroad chromosomes in this triple knockdown condition can be explained by a further reduction of centromere cohesion in the absence of both DDX11 and ESCO2, as compared to single depletions, and at the same time a specific reinforcement of arm cohesion by WAPL knockdown (also see Figure 7).

**Figure 4:**
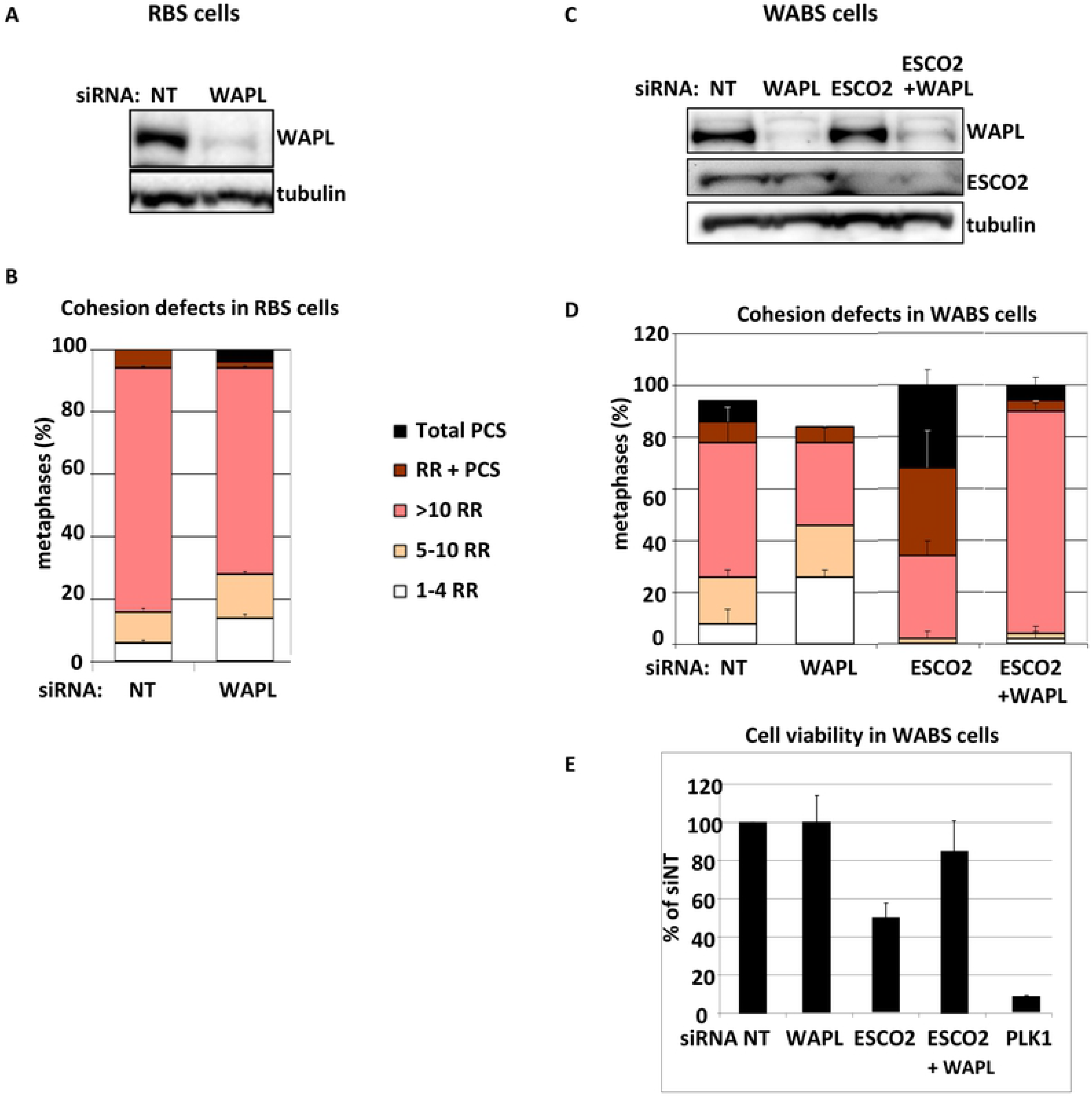
Restoring arm cohesion by WAPL knockdown rescues PCS and lethality. (A,B) RBS cells were transfected with siWAPL and harvested after three days for western blot and analysis of cohesion defects. (C-E) WABS cells were transfected with the indicated siRNAs and harvested after three days for western blot and analysis of SCC. Viability was assessed after four days with a cell-titer blue assay.

In summary, co-depletion of DDX11 and ESCO2 results in severe cohesion loss and lethality, which are both rescued by reinforcing arm cohesion via WAPL knockdown. This indicates that ESCO2 and DDX11 critically facilitate SCC in largely non-overlapping contexts.

### ESCO1 and ESCO2 have separate functions in sister chromatid cohesion

Since ESCO1 and ESCO2 are both to some extent required for SMC3 acetylation, we set out to investigate the extent of their redundancy in SCC by manipulating their expression levels in RBS cells. Depletion of ESCO1 severely aggravated the cohesion defects in RBS cells and we also detected a small effect of ESCO1 depletion in RBS+ESCO2 cells (Figure 5A,B), suggesting that ESCO1 has a role in SCC that cannot be entirely compensated for by ESCO2.

**Figure 5:**
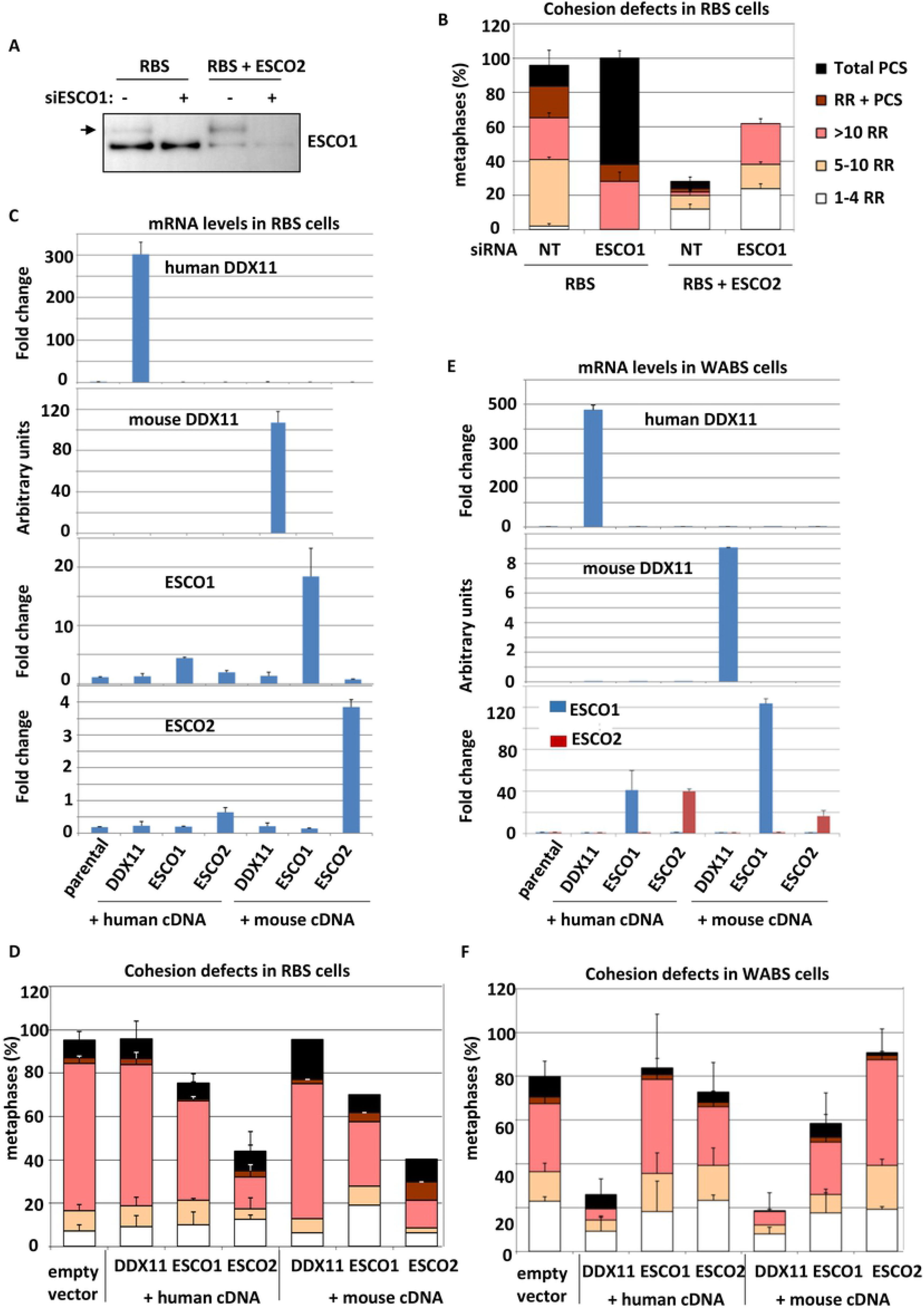
ESCO1, ESCO2 and DDX11 have separable functions in SCC. (A,B) RBS fibroblasts and corrected cells were transfected with siRNA targeting ESCO1 and harvested after three days for Western blot and SCC analysis. Mean and standard deviations of two technical replicates are shown. (C,D) RBS cells were transduced with lentiviral vectors expressing the indicated proteins and selected with puromycin. Overexpression was confirmed with qRT-PCR and cells were analyzed for cohesion defects. Mean and standard deviations of two independent experiments are shown. (E,F) WABS cells were transduced with lentiviral vectors expressing the indicated proteins and selected with puromycin. Overexpression was confirmed with qRT-PCR and cells were analyzed for cohesion defects. Mean and standard deviations of two independent experiments are shown.

Next, we investigated whether overexpression of ESCO1 and ESCO2 could rescue the cohesion defects in RBS cells. To assess possible species-specific effects, we also included their mouse orthologs. Whereas cohesion in RBS could be restored by both human and mouse ESCO2, the effect of ESCO1 was small, indicating that ESCO2 has a unique role in SCC that cannot be performed by ESCO1 (Figure 5C,D). In line with the above described synthetic lethality of ESCO2 and DDX11, DDX11 overexpression could not rescue the cohesion defects in RBS cells. Next, we overexpressed ESCO1, ESCO2 and DDX11 in WABS cells. Cohesion defects in WABS were similarly restored by cDNAs encoding either human or mouse orthologs of DDX11 (Figure 5E,F). Importantly, high levels of either ESCO1 or ESCO2 failed to to compensate for DDX11 inactivation, indicating that cohesion loss resulting from impaired DDX11 function cannot be rescued by increasing SMC3 acetylases. In conclusion, we find that DDX11, ESCO1, and ESCO2 have separable roles in SCC, which are conserved between human and mouse.

### DDX11 deficiency causes reduced DNA replication fork speed

Our findings indicate that ESCO2 and DDX11 act in different pathways leading to SCC establishment. The observed effects of WAPL knockdown suggest that the SCC resulting from the activity of these pathways is at least in part spatially separated at the chromosomes. The failure of ESCO1 and ESCO2 overexpression to compensate for the lack of DDX11 suggests that the activities of these proteins are also mechanistically distinct. DDX11 does not seem to contribute to SMC3 acetylation (Figure 6A-B and Figure S2), although it is possible that fluctuations are masked by SMC3 acetylation that occurs in the context of non-canonical cohesin activities (*e.g.* gene transcription). Indeed, knocking down ESCO2 itself hardly had an effect, and when the ESCO1-dependent effects were eliminated by siRNA, DDX11 overexpression seemed to slightly increase SMC3 acetylation, although this increase was not significant. The interaction of DDX11 with multiple replication fork components [44–46] suggests that it plays a role in cohesin loading in synchrony with DNA replication fork passage. To investigate whether DDX11 also promotes DNA replication itself, we performed a DNA fibre assay. Indeed, we found DDX11 knockdown reduces replication fork speed in RPE1-TERT cells, to levels comparable to those observed after ESCO2 depletion (Figure 6C and 6D). We speculate that DDX11 contributes to SCC via resolving specific secondary DNA structures at the replication fork, thereby promoting cohesin loading, while ESCO2 may connect proper SMC3 acetylation after the fork, to proficient replisome progression by an as yet unclarified mechanism.

**Figure 6:**
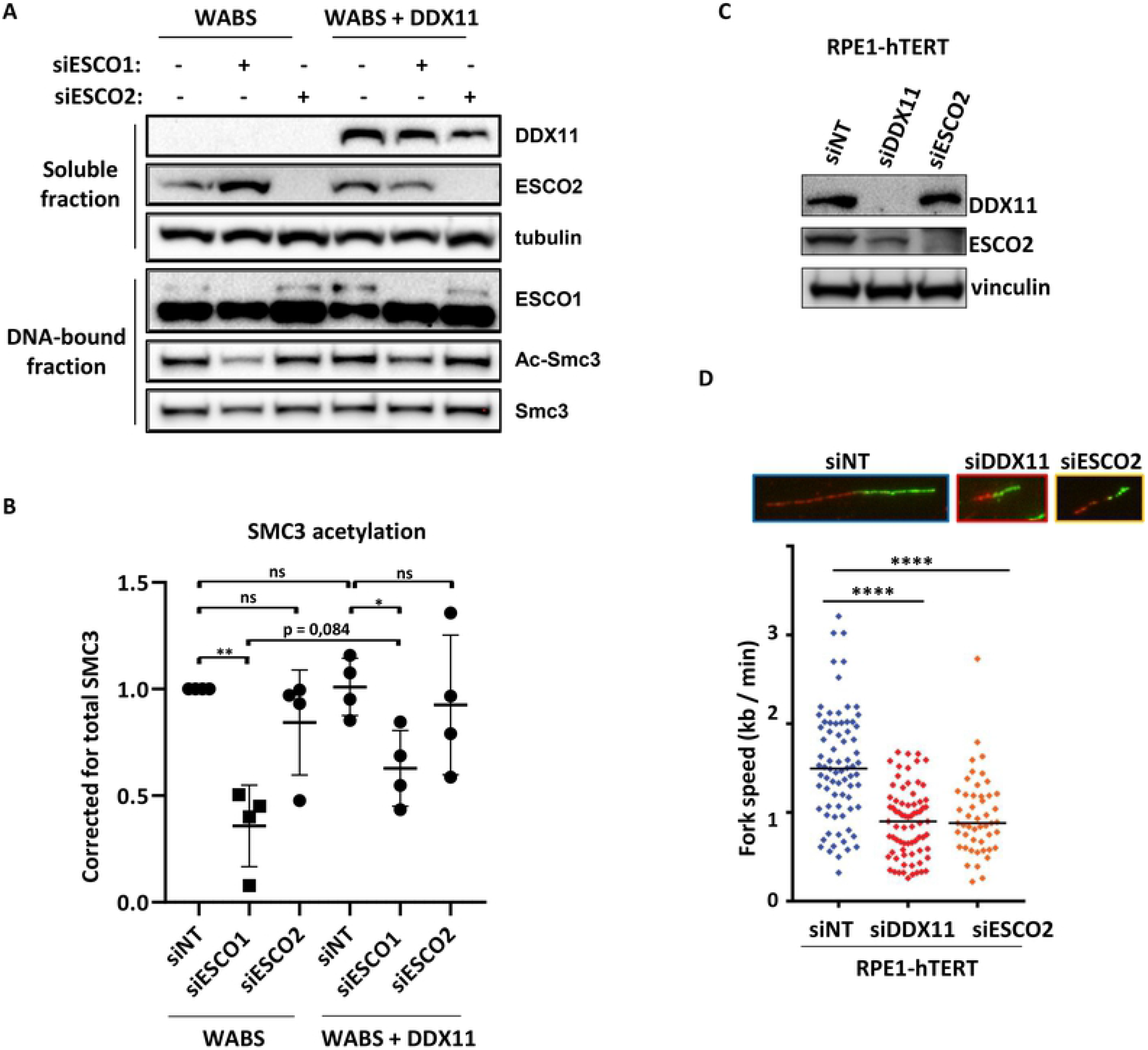
DDX11 deficient cells exhibit DNA replication stress without preventing SMC3 acetylation. (A) Cells were transfected with indicated siRNAs and analyzed by western blot. (B) Quantification of AcSMC3 levels normalized to total SMC3, using image-lab software and. Mean and standard deviations are shown of four independent experiments (provided in Figure S2). P-values were calculated using a two-tailed t-test. * p<0,05; **<0,01; ns not significant. (C,D) RPE1-hTERT cells were transfected with indicated siRNAs for two days, and analyzed by western blot and DNA fiber assay. Replication fork speed of indicated cells was assessed with a DNA fiber assay using a double labeling protocol. Black lines indicate the median. P-values were calculated by a non-parametric one-way ANOVA test. **** p<0,0001. Representative images of DNA fiber tracts are shown.

**Figure 7:**
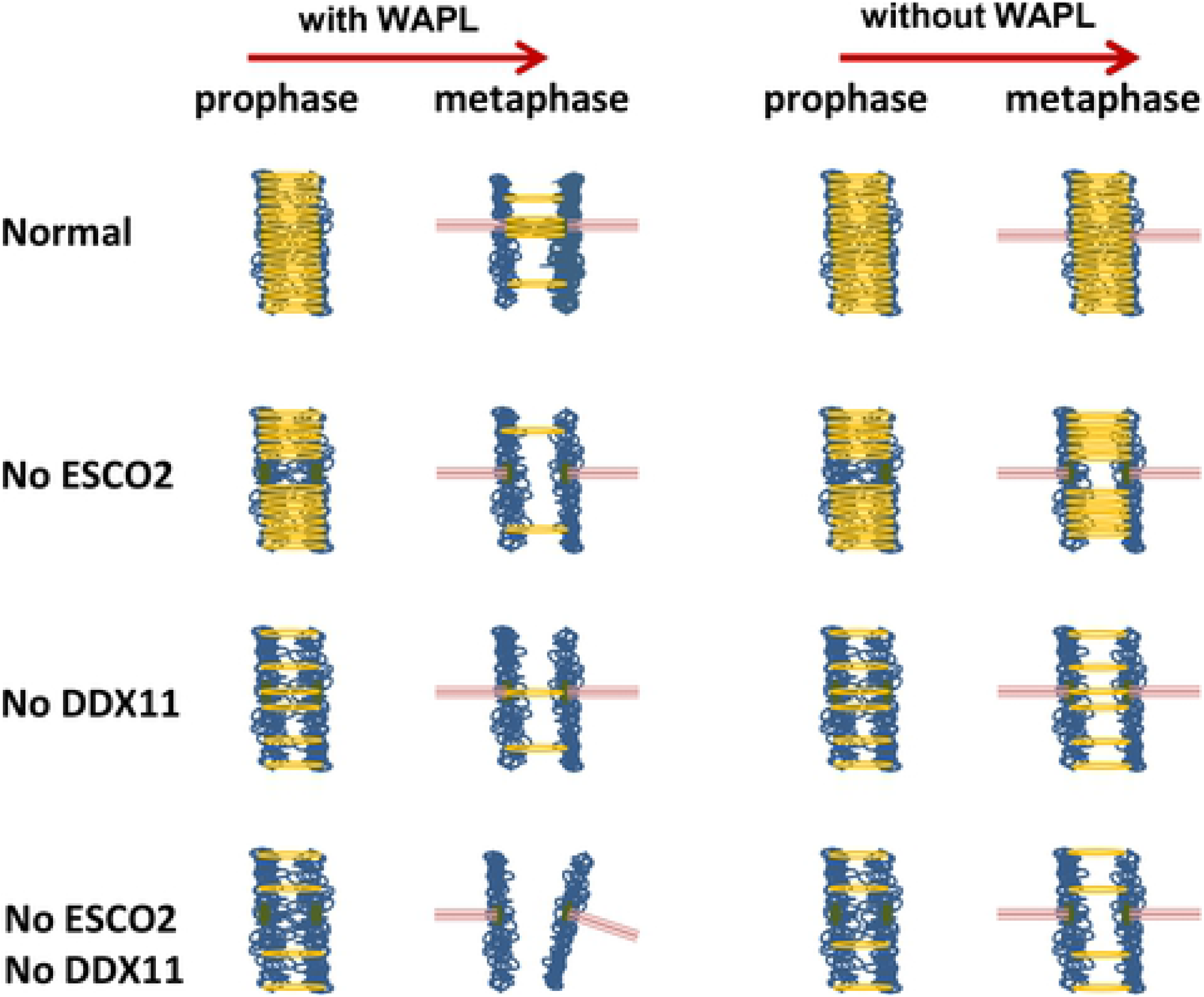
Spatial separation model showing location-specific functions of DDX11, ESCO2 and WAPL in SCC. In normal cells, cohesion is established in S-phase along the entire chromosomes. The bulk of cohesin is removed from chromosome arms during prophase in a WAPL dependent manner, leaving only centromere cohesion intact. The absence of ESCO2 specifically impairs centromere cohesion, which leads to railroad chromosomes that cannot be rescued by WAPL knockdown. DDX11 deficiency, however, affects cohesion on both chromosome arms and centromeres, and WAPL knockdown can partially rescue these effects. Combined loss of ESCO2 and DDX11 depletes cohesin from both arms (DDX11) and centromeres (ESCO2 and DDX11), thereby greatly increasing the chance of PCS. Importantly, in this scenario WAPL depletion prevents PCS, but at the same time increases the number of railroad chromosomes.

## Discussion

In this study we provide evidence of synthetic lethality between ESCO2 and DDX11 in different human cell lines. Lethality correlated with prolonged mitosis and strongly aggravated loss of SCC, suggesting that mitotic arrest is triggered by premature chromatid separation and subsequent mitotic checkpoint activation, as we described before [55]. In line, we also observed signs of apoptosis induction and polyploidy. Importantly, both lethality and cohesion loss were rescued by WAPL knockdown. These observations indicate that ESCO2 and DDX11 promote SCC in largely non-overlapping contexts: loss of only one of these proteins leads to a level of cohesion loss that can still be tolerated, but their combined loss causes near to complete cohesion loss, which is detrimental for cell viability.

An attractive explanation for the synthetic lethality between ESCO2 and DDX11 is that they partially function in spatially separated contexts (Figure 7). A previous study showed that ESCO2 loss particularly affects cohesion in pericentromeric regions [35], which is also illustrated by a large number of railroad chromosomes in RBS cells. It has to be noted that RPE1-ESCO2KO cells also exhibit considerable levels of PCS, indicating that the role of ESCO2 is not entirely restricted to centromeres. WAPL knockdown hardly rescues the cohesion loss in RBS cells, probably because inhibition of WAPL mainly reinforces arm cohesion by blocking the prophase pathway [4, 6, 58]. Indeed, WAPL knockdown partially rescues cohesion defects in DDX11 deficient cells, which occur on both chromosome arms and centromeres. The spatial separation model receives particular support from our observation that in siESCO2 treated WABS cells, WAPL depletion only prevents PCS, but at the same time increases the amount of railroad chromosomes (Figure 3E and Figure 7, bottom right).

The divergence of yeast Eco1 into two vertebrate orthologs, ESCO1 and ESCO2, is accompanied by some functional specialization and differential regulation of their protein levels. For example, ESCO1 is expressed during the whole cell cycle, whereas ESCO2 protein expression peaks in S-phase [35, 36]. This suggests that ESCO2 mainly fulfills the classical role of establishing SCC, whereas ESCO1 regulates cohesins non-canonical activities such as gene transcription. Indeed, ESCO1 was reported to acetylate SMC3 in a replication-independent manner, via a unique interaction with PDS5 [39]. This also explains why we observe a larger effect on SMC3 acetylation of ESCO1 knockdown as compared to ESCO2 knockdown. In line with earlier reports [34, 39, 40], we show here that ESCO1 also contributes to canonical SCC. Although we observed some compensating effects of overexpressing ESCO1 in RBS cells, both knockdown and overexpression studies indicated that ESCO1 and ESCO2 have largely non-overlapping roles in SCC. This might be related to differentially localized fractions of SCC the chromosomes; because ESCO2-dependent SCC preferentially is located at centromeres, ESCO1 may predominantly acetylate cohesin that is present chromosome arms. It will be interesting to further quantify the levels of cohesive cohesin on chromosome arms and centromeres in ESCO1 and ESCO2 deficient human cells. Importantly, we could not distinguish the activities of human or mouse versions of DDX11 or ESCO1/2. Therefore, the paradox that ESCO2 loss is embryonically lethal in mice [59], whereas most human RBS patients contain loss of function mutations in ESCO2 [60], can probably not be explained by differences in the mouse and human ESCO1 and ESCO2 coding sequences. Possibly, the typical acrocentric architecture of murine chromosomes makes them more vulnerable to SCC deprivation and/or ESCO2 deficiency.

We find no clear effect of DDX11 levels on SMC3 acetylation in patient derived cells, although we cannot exclude that DDX11-dependent cohesin loading indirectly contributes to acetylation of cohesin rings that connect newly synthesized sister chromatids. Importantly, we find that increasing SMC3 acetylation, by overexpressing ESCO1 or ESCO2, cannot rescue the cohesion defects in DDX11 deficient cells. Whereas the spatial separation model depicted in Figure 7 may at least partially underlie the severe synthetic lethality between DDX11 and ESCO2, as well as between ESCO1 and ESCO2, it appears insufficient to explain the observations that concern DDX11 and ESCO1. It rather seems that DDX11, ESCO1 and ESCO2 each contribute to different aspects of SCC establishment. DDX11 is thought to resolve certain complex secondary DNA structures including GC-rich regions and/or anti-parallel G-quadruplexes [61, 62]. The duplex or quadruplex resolving capacities of DDX11 helicases may contribute to SCC by facilitating loading of the second strand into cohesin rings, which was reported to require single stranded DNA [63]. Alternatively, in the absence of DDX11, secondary structures that are normally substrates of DDX11 may be prone to breakage, requiring repair and WAPL/PDS5-dependent cohesin removal to provide access of repair factors to the break site (Benedict *et al*, submitted manuscript). Either way, this would inevitably lead to reduced SCC.

## Acknowledgements

We thank Pascal Walther and Mariah Kes for validating DDX11KO and ESCO2KO RPEs, and Hein te Riele for critical reading of the manuscript. This work was supported by the Netherlands Organisation for Scientific research (TOP-GO grant 854.10.013), the Cancer Center Amsterdam (grant CCA2015-5-25) and by the Dutch Cancer Society (Young Investigator grant 10701/2016-2, to JdL). We dedicate this work to the memory of Johan de Winter, who initiated this study.

---

**Figure S1: Deconvolution of siRNAs targeting ESCO2.** WABS fibroblasts and corrected cells were transfected with the indicated siRNAs and cell viability was analyzed after four days, using a cell-titer blue assay.

**Figure S2: DDX11 has no role in SMC3 acetylation.** Four independent experiments in which cells were transfected with indicated siRNAs and analyzed by western blot. Quantifications are shown in Figure 6B.

## Materials and methods

### Cell culture and construction of cell lines

RPE1-hTERT cells (American tissue culture collection) and SV40 transformed fibroblasts, including WABS (Van der lelij et al., 2010), RBS [36] and LN9SV control (Hermsen MA 1996), were cultured in Dulbecco’s Modified Eagles Medium (DMEM, Gibco), supplemented with 10% FCS, 1 mM sodium pyruvate and antibiotics.

CRISPR-Cas9 was used to construct DDX11 and ESCO2 knockouts in RPE1 cells. The generation of RPE1-hTERT_TetOn-Cas9_TP53KO cells is also described in a currently submitted manuscript (Benedict *et al*.). Briefly, Cas9 cDNA was cloned into the pLVX-Tre3G plasmid (Clontech) and lentiviral Tre3G-Cas9 and Tet3G particles were produced in HEK293T cells using the Lenti-X HT packaging system (Clontech). Transduced cells were selected with 10 μg/mL puromycin and 400 μg/mL G418. Cells were treated with 100 ng/mL doxycycline (Sigma-Aldrich) to induce Cas9 expression and transfected with 10 nM synthetic crRNA and tracrRNA (Dharmacon or IDT) using RNAiMAX (Invitrogen). The following crRNA sequences were used: TP53 (CCATTGTTCAATATCGTCCG), DDX11-specific (GGCTGGTCTCCCTTGGCTCC), ESCO2 (TAAGTGGTACCTCAATCCAC). Single clones were assessed by Sanger sequencing using the following primers: TP53-Fw (GAGACCTGTGGGAAGCGAAA, TP53-Rv GCTGCCCTGGTAGGTTTTCT), DDX11-Fw (AACAACCCACCCTCCCCAAG), DDX11-Rv (TGCCTCACTCTCTCCAGACC), ESCO2-Fw (ATCAAAAAGGTAGAAGATGTCCAAGAAC), ESCO2-Rv (GCCTGTTTGATGGGTTCTGC).

### Proliferation assays

Adherent cells, the IncuCyte Zoom instrument (Essen Bioscience) was used. RPE1 cells (1500 / well) and fibroblasts (3000 / well) were seeded in 96-wells plates and imaged every 4 hours with a 10x objective. IncuCyte software was used to quantify confluence from four non-overlapping bright field images per well, for at least three replicate wells. Doubling time was calculated for the period required to grow from approximately 30% to 70% confluence, using the formula doubling time (h) = required time (h) * log(2) / (log(confluence endpoint(%)) – log(confluence starting point(%))).

### siRNA transfections

For knockdown experiments, 25 nM siRNA (Dharmacon) was transfected using RNAiMAX (Invtrogen). Sequences: non-targeting siRNA UAAGGCUAUGAAGAGAUAC, siDDX11 GCAGAGCUGUACCGGGUUU, CGGCAGAACCUUUGUGUAA, GAGGAAGAACACAUAACUA, UGUUCAAGGUGCAGCGAUA, siESCO1 GGACAAAGCUACAUGAUAG, siESCO2 CAAAAUCGAGUGAUCUAUA GAGAGUAGUUGGGUGUUUA, AAUCAAGGCUCACCAUUUA, GAAGAAAGAACGUGUAGUA, siUBB CCCAGUGACACCAUCGAAA, GACCAUCACUCUGGAGGUG, GUAUGCAGAUCUUCGUGAA, GCCGUACUCUUUCUGACUA.

### Proliferation assays

Proliferation assays were performed in 96-wells plates. Cells were counted and seeded in at least triplicates in a total volume of 100 μl medium. Optimized cell densities were: WABS cells 3,000/well, RBS cells 4,000/well, RPE1 cells 1500/well. For WABS and RBS cells, cells were incubated with 10 μl CellTiter-Blue reagent (Promega) for 2–4 h and fluorescence (560_Ex_/590_Em_) was measured in a microplate reader (TriStar LB 941, Berthold Technologies). To monitor cell growth of RPE1 cells, the IncuCyte Zoom instrument (Essen Bioscience) was used. Cells were imaged every 4h with default software settings and a 10x objective. The IncuCyte software was used to quantify confluence from four non-overlapping bright field images.

### qRT-PCR

Total RNA was extracted with the High Pure Isolation Kit (Roche) and cDNA was prepared with the iScript cDNA Synthesis Kit (Biorad). Quantitative reverse transcription polymerase chain reaction (qRT-PCR) was performed using SYBR Green (Roche) on a LightCycler 480 (Roche). Levels were normalized to the geometric mean of at least two housekeeping genes. Primer sequences: hDDX11-Fw AACCTGTTCAAGGTGCAGCGATAC, hDDX11-Rv GAGAAGCTGGTCGCAGGGT mDDX11-Fw TTGTGGCTGTTTTGGGAGGTAATG, mDDX11-Rv CACCTGGCTCTGAAAGAGAAAGTC h/mESCO1-Fw CCTGGTGCTGCTCAACATT, h/mESCO1-Rv CAGGAGTGGGATCTGAGAAAGC m/hESCO2-Fw ATCAAAAAGGTAGAAGATGTCCAAGAAC, m/hESCO2-Rs GCCTGTTTGATGGGTTCTGC HPRT1-Fw TGACACTGGGAAAACAATGCA, HPRT-Rv GGTCCTTTTCACCAGCAAGCT TBP-Fw TGCACAGGAGCCAAGAGTGAA, TBP-Rv CACATCACAGCTCCCCACCA B2M-Fw ATGAGTATGCCTGCCGTGTGA, B2M-Rv GGCATCTTCAAACCTCCATG

### Immunoblotting

Cells were lysed in lysis buffer (50 mM Tris-HCl pH 7.4, 150 mM NaCl, 1% Triton X-100) with protease-and phosphatase inhibitors (Roche). For DNA-bound protein fractions, cells were lysed in lysis buffer for 10 min and centrifuged at 1300 g for 10 min. The pellet was subsequently lysed in lysis buffer containing 5 units/uL benzonase nuclease (Sigma) for 1 h and centrifuged at maximum speed for 5 min. Proteins were separated by 3-8% or 8-16% SDS-PAGE (NU-PAGE or BioRad) and transferred to immobilon-P membranes (Millipore). Membranes were blocked in 5% dry milk in TBST-T (10 mM Tris-HCl pH 7.4, 150 mM NaCl, 0.04% Tween-20), incubated with primary and peroxidase-conjugated secondary antibodies (DAKO Glostrup, Denmark) and bands were visualized by chemoluminescence (Amersham). Antibodies used for detection are mouse anti-DDX11 (B01P, Abnova), mouse anti-α-tubulin (B-5-1-2, Santa Cruz #sc-23948), mouse anti-vinculin (H-10, Santa Cruz sc-25336), guinea pig anti-ESCO2 [36], mouse anti-ESCO1 (gift from JM Peters), rabbit anti-PARP (9542, Cell signaling), AcSM3 (gift from K Shirahige), rabbit anti-SMC3 (A300-060A, Bethyl), rabbit anti-WAPL (A300-268, Bethyl), mouse anti-vinculin (H-10, sc-25336, Santa Cruz).

### Flow cytometry

Cells were harvested and washed in PBS and fixed in ice-cold 70% EtOH. For mitosis detection, cells were incubated with rabbit anti-pS10-Histone H3 (Millipore) for 1 h and with Alexa Fluor 488 goat-anti-rabbit (Invitrogen) for 30 min. Cells were washed and resuspended in PBS with 1:10 PI/RNase staining buffer (BD Biosciences) and analysed by flow cytometry on a BD FACSCalibur (BD Biosciences). Cell cycle analysis was conducted with BD CellQuest software (BD Biosciences).

### Analysis of cohesion defects

Cells were incubated with 200 ng/mL demecolcin (Sigma-Aldrich) for 20 minutes. Cells were harvested, resuspended in 0.075 M KCl for 20 minutes and fixed in methanol/acetic acid (3:1). Cells were washed in fixative three times, dropped onto glass slides and stained with 5% Giemsa (Merck). Cohesion defects were counted in 25 metaphases per slide on two coded slides per condition.

### DNA Fiber analysis

Cells were pulse-labeled with 25 μM Chlorodeoxyuridine (CldU) for 20 minutes, followed by 20 minutes 250 μM Iododeoxyuridine (IdU). Approximately 3000 cells were lysed in 7 μL spreading buffer (200 mM Tris-HCl PH 7.4, 50 mM EDTA and 0.5% SDS). Fibers were spread on Superfrost microscope slides, which were tilted ~15°, air-dried for several minutes and fixed in methanol:acetic acid (3:1). DNA was denatured with 2.5 M HCl for 75 minutes, blocked in PBS + 1% BSA + 0.1% Tween20 and incubated for 1 hour with rat-anti BrdU (1:500, Clone BU1/75, Novus Biologicals) and mouse-anti-BrdU (1:750, Clone B44, Becton Dickinson). Slides were fixed with 4% paraformaldehyde for 10 minutes, incubated for 1.5 h with goat-anti-mouse Alexa 488 and goat-anti-rat Alexa 594(both 1:500, Life technologies) and mounted with Vectashield medium. Images of DNA fibers were taken with a Zeiss AxioObserver Z1 inverted microscope using a 63x objective equipped with a Hamamatsu ORCA AG Black and White CCD camera. Fiber tract lengths were assessed with ImageJ and μm values were converted into kilobases using the conversion factor 1 μm = 2.59 kb [64, 65].

